# Near-field potentials index local neural computations more accurately than population spiking

**DOI:** 10.1101/2023.05.11.540026

**Authors:** David A. Tovar, Jacob A. Westerberg, Michele A. Cox, Kacie Dougherty, Mark T. Wallace, André M. Bastos, Alexander Maier

## Abstract

Local field potentials (LFP) are low-frequency extracellular voltage fluctuations thought to primarily arise from synaptic activity. However, unlike highly localized neuronal spiking, LFP is spatially less specific. LFP measured at one location is not entirely generated there due to far-field contributions that are passively conducted across volumes of neural tissue. We sought to quantify how much information within the locally generated, near-field low-frequency activity (nfLFP) is masked by volume-conducted far-field signals. To do so, we measured laminar neural activity in primary visual cortex (V1) of monkeys viewing sequences of multifeatured stimuli. We compared information content of regular LFP and nfLFP that was mathematically stripped of volume-conducted far-field contributions. Information content was estimated by decoding stimulus properties from neural responses via spatiotemporal multivariate pattern analysis. Volume-conducted information differed from locally generated information in two important ways: (1) for stimulus features relevant to V1 processing (orientation and eye-of-origin), nfLFP contained more information. (2) in contrast, the volume-conducted signal was more informative regarding temporal context (relative stimulus position in a sequence), a signal likely to be coming from elsewhere. Moreover, LFP and nfLFP differed both spectrally as well as spatially, urging caution regarding the interpretations of individual frequency bands and/or laminar patterns of LFP. Most importantly, we found that population spiking of local neurons was less informative than either the LFP or nfLFP, with nfLFP containing most of the relevant information regarding local stimulus processing. These findings suggest that the optimal way to read out local computational processing from neural activity is to decode the local contributions to LFP, with significant information loss hampering both regular LFP and local spiking.

**Author’s Contributions:** Conceptualization, D.A.T., J.A.W, and A.M.; Data Collection, J.A.W., M.A.C., K.D.; Formal Analysis, D.A.T. and J.A.W.; Data Visualization, D.A.T. and J.A.W.; Original Draft, D.A.T., J.A.W., and A.M.; Revisions and Final Draft, D.A.T., J.A.W., M.A.C., K.D., M.T.W., A.M.B., and A.M.

**Competing Interests:** The authors declare no conflicts of interest.

## Introduction

The local field potential (LFP) is a complex intracranial neural signal comprising transmembrane potentials arising from incoming synaptic inputs (Buzsáki et al., 2012; Mitzdorf, 1985), sodium currents (Ray et al., 2008), calcium currents (Schiller et al., 2000), and gap junctions (Traub and Bibbig, 2000). LFP captures graded potentials in addition to the all-or-none response of the action potential, effectively providing a broader population response (Bijanzadeh et al., 2018). As a result, the information contained in LFPs often complements what is found from action potentials (Leszczyński et al., 2020; Mineault et al., 2013), and in some cases explains behavioral responses better than action potentials (Pesaran et al., 2002). Additionally, LFPs show more consistency across recording sessions than population spiking activity, as they are not affected as much by the position of the electrode relative to the recorded population (Bédard et al., 2004). This stability can be partially explained by the LFPs proclivity to diffuse through space via passive volume conduction. However, this otherwise helpful property can be problematic for identifying the structures involved in generating or receiving neural signals. Volume conducted signals originate at cortical columns immediately surrounding the electrodes, as well as structures far outside the structure of interest (Bertone-Cueto et al., 2020; Kajikawa et al., 2017; Kajikawa and Schroeder, 2011). There are a number of known instances where a brain structure thought to have been involved in producing a neural signal was in fact the result of volume conducted LFPs from nearby structures (Bertone-Cueto et al., 2020; Kajikawa et al., 2017; Lalla et al., 2017). Accounting for this volume conduction is important given that stimulus features are processed preferentially across sensory cortical areas, within areal maps comprised of columns, and even across the layers of a cortical column (Tovar et al., 2020).

Reports of the extent of volume conduction through neural tissue range from a few hundred micrometers (Katzner et al., 2009; Xing et al., 2009) to several centimeters (Kajikawa and Schroeder, 2015, 2011; Kreiman et al., 2006; Nauhaus et al., 2009). A number of factors contribute to this variability. Cell morphology is a considerable factor, with modeling showing that pyramidal cells, due to their asymmetry, give rise to larger LFP spread than any other cell type (Lindén et al., 2011). The spontaneous correlation between cells at rest as well as during stimulus presentation is also a factor, and is affected by features such as the presence of horizontal cells and the particular stimulus features encoded, which can affect passive spread by an order of magnitude (Leski et al., 2013; Lindén et al., 2011; Rosenbaum et al., 2017). Lastly, there is volume conducted passive spread when neural activity in one brain area elicits activity in neighboring brain areas, propagating as a traveling wave (Sato et al., 2012; Zanos et al., 2015). These factors can make precise spatial localization of the neural generators of LFP rather challenging. Notably, the extent of the volume conducted component can be investigated through careful transformation of the LFP signal. This is possible by collecting LFP across an area or volume of cortex, such as individual cortical columns. These LFP measures can then be transformed by computing the second spatial derivative of the LFP (Mitzdorf, 1985), known as current source density, or CSD. The CSD estimates the (mostly synaptic) currents sinks and sources underlying the near-field LFP, largely eliminating volume-conducted components. CSD is a complex signal, requiring more principle components than LFP and population spiking to explain signal variance (Einevoll et al., 2007; Schaefer et al., 2017). Importantly, the CSD can be re-summed into an estimate of locally generated, near-field LFP (nfLFP – at the columnar microcircuit scale) with minimal contamination by volume conduction, previously also described as LFP_cal_ (Kajikawa and Schroeder, 2011; Westerberg et al., 2022, 2021). However, investigation using the nfLFP signal is limited. The nfLFP has primarily been used to quantify the amount of volume conduction in the original LFP signal (Kajikawa and Schroeder, 2011). In contrast, this more local signal has not been used to study how LFP information content is affected by volume conduction.

Here, we used multivariate pattern analysis to study the information content of volume conduction-containing versus locally generated LFP in primate primary visual cortex (V1) during visual presentation of multifeatured stimuli. Since certain stimulus features are uniquely processed across brain areas and thus might yield different degrees of volume conduction, we studied stimulus features matching V1 response preferences as well as stimulus features that evoke heightened responses outside of V1 (i.e., in higher visual cortex). We find that decoding performance for features primarily processed within V1 (such as orientation, ocular dominance) suffer from volume conduction effects while decoding performance for stimulus features primarily processed outside of V1 (such as stimulus sequences) appear enhanced by volume conduction. These findings suggest that the bulk of volume conducted signals in V1 originates in higher visual cortex and does not contribute to local processing. Thus, LFP can be less rather than more informative than near-field signals. At the same time, we found that local population spiking can be less informative than near-field signals, suggesting that certain stimulus-relevant information remains subthreshold, but can be recovered by decoding near field LFP instead of spiking. Additionally, we found that volume conduction differentially affected information content across LFP frequency bands associated with feedforward versus feedback processing (Bastos et al., 2015, 2012; Belitski et al., 2008; Peter et al., 2019; Van Kerkoerle et al., 2014). Our findings demonstrate that volume conducted signals frequently mask information in locally generated low-frequency signals. Thus, computational elimination of volume-conducted contributions transforms LFP into the most informative neural readout within cortical microcircuitry, largely exceeding the informative value of population spiking responses (which in turn exceeds the informative value of simultaneously collected single neurons (Trautmann et al., 2019)).

## Methods

### Animal care and surgical procedures

Procedures were in accordance with National Institutes of Health Guidelines, Association for Assessment and Accreditation of Laboratory Animal Care Guide for the Care and Use of Laboratory Animals, and approved by the Vanderbilt Institutional Animal Care and Use Committee following United States Department of Agriculture and Public Health Services policies. Two macaque monkeys (*Macaca radiata*: monkey E48 [male], monkey I34 [female]) underwent a series of surgeries implanting MR compatible head posts and cranial recording chambers positioned over one hemisphere of V1. A craniotomy was performed concurrent with the location of the recording chamber. All surgical procedures were performed under general anesthesia. Anesthetic induction was performed with ketamine (5-25 mg/kg). Monkeys were then catheterized and intubated. Surgeries were performed under aseptic conditions. N_2_O/O_2_, isoflurane (1-5%) anesthesia was used. Vital signs were monitored continuously. Postoperative antibiotics and analgesics were administered. Additional descriptions of animal care and surgical procedures can be found elsewhere (Westerberg et al., 2020a, 2020b, 2019).

### Magnetic resonance imaging

Magnetic resonance (MR) imaging was used to guide recording chamber implant surgeries as well as to guide linear electrode array penetrations. All MR scans were conducted with animals under general anesthesia per the procedures described in *Animal care and surgical procedures*. Scans were obtained using a Philips 3T MR scanner. T1-weighted 3D MPRAGE scans were acquired with a 32-channel head coil equipped for sense imaging. Images were acquired using 0.5 mm isotropic voxel resolution with the following parameters: repetition 5 s, echo 2.5 ms, and flip angle 7°.

### Visual display and stimuli

Monkeys viewed stimuli presented on a 20” CRT monitor at 60 or 85 Hz. Stimuli were presented through a custom mirror stereoscope allowing for monocular or binocular presentation of stimuli (Figure 1A) (Cox et al., 2019b; Dougherty et al., 2019, 2021). Prior to performance of the main task, monkeys performed a stereoscope calibration task as to eliminate any potential confound of binocular disparity (see Tovar and Westerberg et al., 2020). Each recording session (N = 61, monkey E48: 48, monkey I34: 13) comprised several hundred trials each with sequences of 1-5 stimulus presentations (Figure 1B). Stimulus displays were generated using MonkeyLogic (Asaad and Eskandar, 2008; Hwang et al., 2019). Monkeys initiated trials by fixating within 1 degree of visual angle (dva) of a central fixation dot. Following a short (300 ms) fixation period, stimuli appeared in sequence (1-5 stimuli) each for 200 ms with a 200 ms inter-stimulus interval. If monkeys maintained fixation throughout the stimulus sequence, they received a juice reward before an inter-trial interval ensued. If monkeys failed to maintain fixation throughout the trial, they received a brief timeout before the inter-trial interval. Each stimulus presented was a sinusoidal bar grating stimulus localized at the visual receptive field of the cortical column (see *Receptive field mapping* below). The stimuli maintained the same size, spatial frequency, and phase throughout each recording session (optimized for the column multi-unit activity, see Cox et al., 2013, 2019a, 2019b), but had variable orientation and eye-of-origin across trials. Stimuli were presented monocularly.

**Figure 1:**
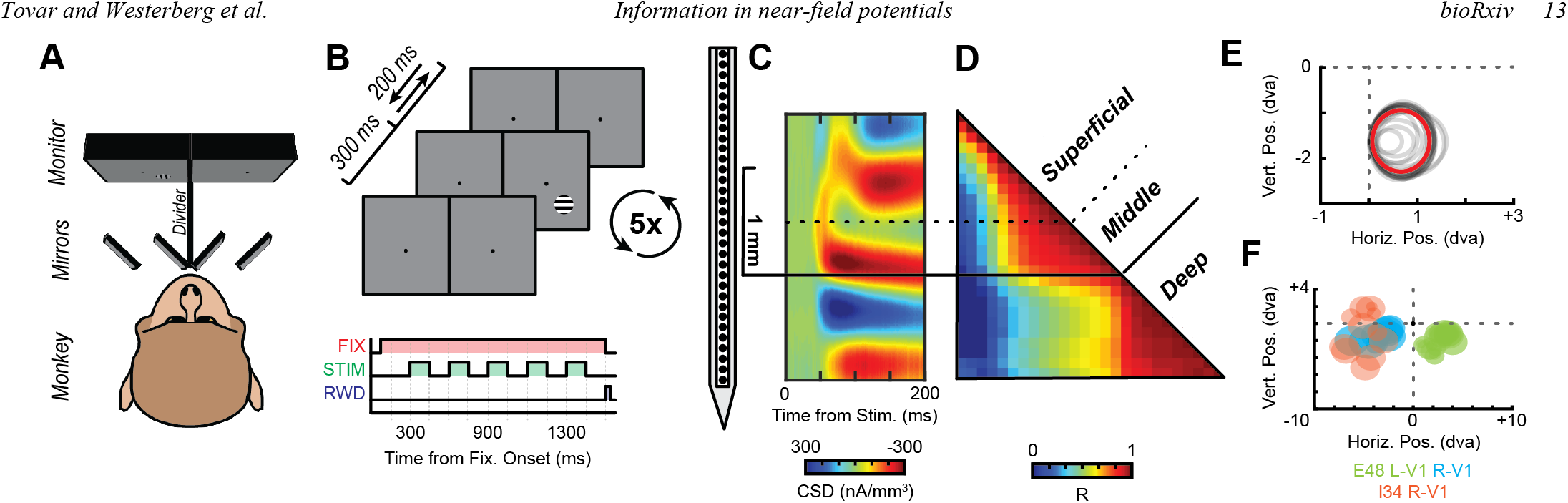
Experimental setup and cortical laminar alignment of linear multielectrode array. A. Stereoscope setup for stimulus presentation. Monkey is positioned behind a series of mirrors which provide visual stimulation to each of the eyes independently. B. Stimulus presentation sequence. Monkeys fixated for 300 ms after which a stimulus was presented to the receptive field of the V1 column for 200 ms. 5 stimuli were presented sequentially with 200 ms inter-stimulus intervals while monkeys maintained central fixation. C. Laminar electrode alignment across sessions using current source density (CSD). For each session (n=61), CSD was computed, and the L4/5 boundary identified as the earliest stimulus evoked net depolarization (current sink) with an accompanying deeper source. Color plot shows session average CSD response to stimuli in the columnar population receptive field (RF) following alignment. D. LFP coherence was also used to confirm laminar alignment. Color plot shows spatial profile of contact-by-contact correlations across time (512 ms moving window) during each session, averaged across all sessions following laminar alignment. E. Representative RF mapping for a single session. Each gray circle represents the estimated multiunit RF for a recording site identified to be in cortex. The red circle indicates the average along contacts. All circles overlap, indicating perpendicular penetration. F. Average columnar RF for each session where color indicates the monkey-hemisphere combination where the column was recorded.

### Neurophysiological procedure

Broadband (0.5-12.207 kHz) neurophysiological signals were recorded during task performance. Signals were amplified and digitized at 30 kHz using a 128-channel Cerebus Neural Signal Processing System (Blackrock Microsystems). LFP signals were downsampled to 1 kHz. All neural recordings were performed using 24-or 32-channel linear microelectrode arrays with 0.1 mm interelectrode spacing (S-probe, U-Probe, V-Probe – Plexon; Vector array – NeuroNexus) positioned orthogonal to the cortical surface in dorsal V1. Microelectrode recording contacts had impedances between 0.2-0.8 Mohms. Electrode arrays were held in position using a custom Narishige micromanipulator. Electrode arrays were interfaced with the amplifier system using the Blackrock analog head stage. Gaze was measured binocularly at 1 kHz using an Eyelink system (SensoMotoric Instruments). All recordings took place in a radio frequency-isolated booth.

### Population spiking (MUA)

We used multiunit spiking activity (MUA) to estimate neural population dynamics (Trautmann et al., 2019). A full description of the procedure to extract MUA can be found at (Legatt et al., 1980; Logothetis et al., 2001; Self et al., 2013; Shapcott et al., 2016; Supèr and Roelfsema, 2005; Teeuwen et al., 2021; Tovar et al., 2020; Westerberg et al., 2020). Briefly, broadband extracellular voltage time series were filtered between 0.5-5 kHz, which matches the spectral peak of spiking activity. The filtered signal was full wave rectified to obtain power. Lastly, the signal was low pass filtered at 0.25 kHz. For filtering, we used a 4^th^-order bidirectional Butterworth filter.

### Receptive field mapping

Receptive field mapping was performed at the beginning of the neural recordings to ascertain the receptive field of the cortical column being recorded from as well as to confirm an orthogonal electrode array penetration into V1 (Figure 1E-F). Multiunit and LFP activity was measured while a series of stimuli were presented in the contralateral lower quadrant of the visual hemifield relative to the position of the recording chamber. Monkeys fixated a central fixation dot while 1-5 stimuli were presented.

Successful maintenance of fixation throughout the stimuli presentations yielded a juice reward. Qualitative (auditory evaluation) and quantitative assessment of the visual responses was performed online. More detailed description of the procedure is detailed elsewhere (Cox et al., 2013). Monkeys proceeded to perform the main task described in *Visual display and stimuli* if there was an observable receptive field which was consistent along cortical depth, indicative of an orthogonal presentation. The measured receptive field in this task was used as the location for stimuli presentation in the main task.

### Current source density and laminar alignment

Current source density (CSD) served to identify the location of the electrode relative to the layers of V1 (Mitzdorf, 1985). The spatiotemporal profile of CSD has a distinct pattern which allows for the reliable identification of the boundary between the granular input layers and the infragranular layers of V1 (Schroeder et al., 1998). To compute the CSD from the LFP, we used a previously describe procedure (Nicholson and Freeman, 1975):

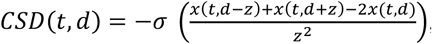

where the CSD at timepoint *t* and at cortical depth *d* is the sum of voltages *x* at electrodes immediately above and below (*z* is the interelectrode distance) minus 2 times the voltage at *d* divided by the interelectrode-distance-squared. That computation yields the voltage local to *d*. To transform the voltage to current, we multiplied the result by *−*σ, where σ is a previously reported estimate of the conductivity of cortex (Logothetis et al., 2007). In addition to using CSD for laminar alignment (Figure 1C), we confirmed positioning of the electrode array relative to V1 layers by identifying reliable patterns in the correlations between LFP across electrodes (Figure 1D) and in the LFP power spectral density (PSD, Figure 2) through previously reported means (Maier et al., 2010; Westerberg et al., 2019).

**Figure 2:**
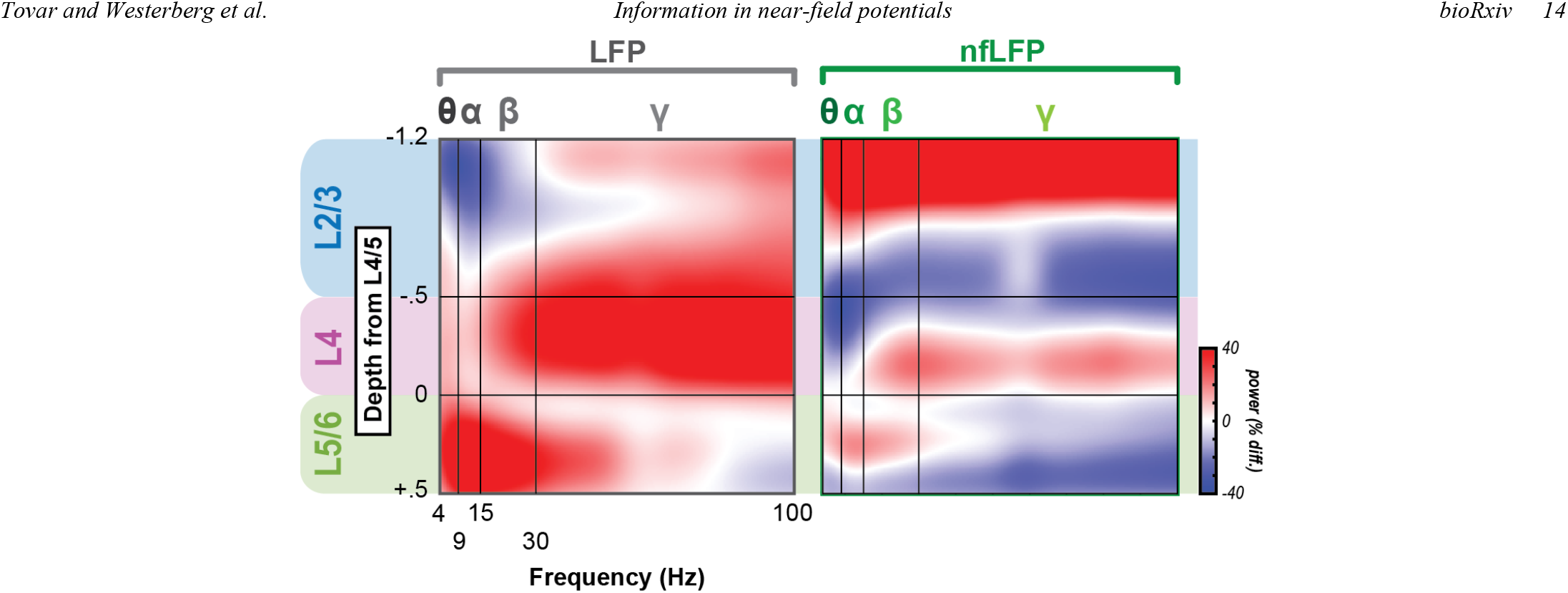
Laminar power spectral density (PSD) for regular, volume conduction polluted LFP and computationally derived, locally generated nfLFP. Left column shows regular LFP and right column, the nfLFP. PSD was normalized by finding the average power for each frequency along cortical depth. Then, power for each electrode array contact for each frequency was taken as the percent difference from the column average. Profiles were first normalized at the session level, then averaged across sessions (n=61). Ordinates are cortical depth relative to the L4/5 boundary. Abscissa represents frequency. Red indicates greater than column average power for that frequency at that depth. Blue indicates lower than column average power. The transformation of LFP to nfLFP modifies the laminar spectral profile of field potential power, with the main motif (i.e., gamma power peaking in the granular layers and lower frequencies dominating lower layers) remaining.

### Locally generated (near-field) LFP recalculation

The configuration of microelectrodes in a linear array provides the opportunity to recalculate the low-frequency LFP signal with minimal contamination by volume-conduction. We calculated the locally generated component of the LFP from the measured laminar CSD (nfLFP) using a previously described model (Kajikawa and Schroeder, 2011; Nicholson and Llinas, 1971):

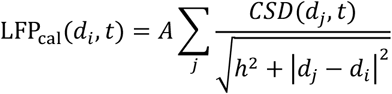

where LFP_cal_ *a*t depth *i (d*_*i*._) for each timepoint *t* is taken as the sum of CSD at depths *j (d*_*j*_) for each timepoint divided by the Euclidean distance to account for the attenuating impact of local currents on distant field potentials. The factor *A* acts only as a scaling factor. Since we cannot accurately estimate the magnitude of the one-dimensional CSD-derived waveform, this parameter was set to 1. This omission is consistent with previous reports (Kajikawa and Schroeder, 2011) and limits our comparisons of volume-conducted LFP and the locally generated nfLFP to the shape of the waveforms. However, magnitude differences can be observed between conditions for the volume-conducted and locally generated LFP, independently (Figure 3). Also, for our purposes, we set *h t*o 0 as we assume that the observed CSD and recalculated LFP are colocalized.

**Figure 3:**
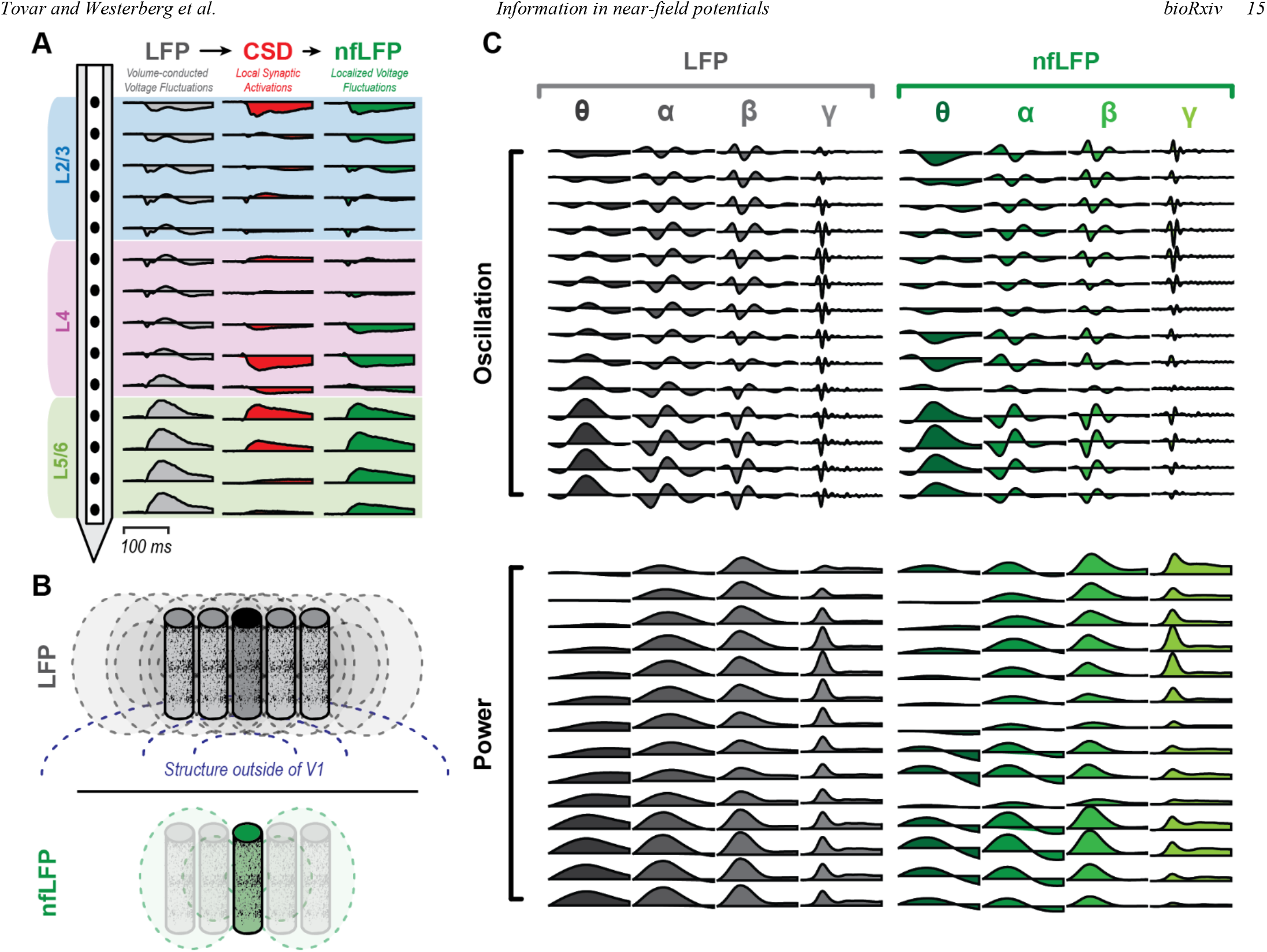
Calculation of locally generated nfLFP from regular, volume-conduction polluted LFP for a representative session. A. Procedure for deriving nfLFP. Regular LFP is recorded at each linear electrode array contact. CSD is computed as the second spatial derivative of the LFP signal. nfLFP is then calculated from the CSD as the sum of field potentials generated at each of the electrodes, accounting for attenuation of magnitude as a function of distance. Blue, purple, and green backgrounds indicate laminar compartments. B. Cartoon exemplifying the concept behind the nfLFP procedure. Cylinders represent cortical columns. Dashed lines represent field potentials. LFP signals stem from both locally generated and volume-conducted signals, including LFP generated in nearby cortical columns as well as deeper neural structures. The nfLFP procedure attenuates or eliminates signals generated outside of the recorded column to estimate the LFP that is locally generated. C. Spectral decomposition into frequency bands for both regular LFP signal as well as nfLFP.

### LFP frequency analysis

Once the volume conduction-containing LFP and the recalculated, locally generated nfLFP were isolated, we performed a filtering step (where necessary) to investigate differences that might exist with respect to component frequency bands. Filtering was done using a bidirectional bandpass 4^th^ order Butterworth filter (Maier et al., 2011). Filtering was performed on the raw neurophysiological signal prior to extracting trials. All power spectral density analyses were calculated on both raw and recalculated neurophysiological signal through a Fourier transform with a 512 ms window and no window overlap. Data were not event-locked for PSD analyses, but averaged across the entire experiment duration instead.

### Multivariate pattern analysis

To track sensory information flow within the laminar microcircuit, we applied multivariate pattern analysis (MVPA) using CoSMoMVPA (Oosterhof et al., 2016). To do so, we assembled two-dimensional neuronal response matrices (NRMs) that contained the millisecond-by-millisecond population neural (LFP, nfLFP or MUA) response at each electrode array contact/channel as a function of stimulus presentation. Each repetition of a stimulus presentation elicits a different response across each of the dimensions (representing electrode contacts) of the NRM. We extracted information regarding (i.e., decoded) grating orientation, the eye that the stimuli were presented to (eye-of-origin) and the relative position of each stimulus within the stimulation sequence using a linear discriminant analysis (LDA) classifier trained and tested for each session. Three quarters of the data was used to train an MVPA classifier. The remaining ¼ of the NRMs were used to determine decoding performance. Training subsets and testing subsets were sub-sampled to ensure the same number of trials per stimulus condition. Decoding accuracy was defined as the number of trials over the total number of trials that the classifier was able to correctly identify for each session. We performed this computation on a millisecond-by-millisecond basis within each recording session to obtain classifier performance as a function of time (time-resolved MVPA). The resulting time courses of decoding accuracy were compared to a randomized shuffle control to determine statistical significance. To correct for multiple comparisons, we used false discovery rate (FDR) adjusted p-values with α = 0.001. This analysis was performed on the LFP, nfLFP, and MUA signals for electrodes contained within each laminar compartment.

To compare how stimulus features differed and evolved over time and space, we used a searchlight analysis on all three signals (Etzel et al., 2013; Tovar et al., 2020). For this analysis, a linear discriminant analysis (LDA) classifier was trained and tested for each session using one electrode and its immediate neighboring electrodes at each time ranging from 100 ms prior to stimulus presentation to 400 ms after stimulus onset. For signal differences, significance was evaluated against zero. In addition to the broadband LFP and nfLFP, a searchlight analysis was also performed on the frequency filtered nfLFP and LFP signals. All searchlight maps were thresholded using Wilcoxon signed rank tests that evaluated for statistical significance compared to chance decoding, with FDR correction. Chance decoding is defined as the percent correct obtained when the classifier guesses randomly.

To understand how stimulus features are shared across signal frequencies throughout a stimulus presentation, we devised a novel frequency generalization analysis The idea behind the frequency generalization matrix was inspired by the time generalization analysis used in a number of previous studies (Carlson et al., 2011; King Dehaene and S., 2014; Tovar et al., 2020). After construing time-frequency spectrogram (described above), we trained an LDA classifier at one LFP frequency band and then tested the classifier at all remaining frequency bands to reveal shared stimulus information across frequency bins. We iteratively repeated this process until we obtained a matrix of all frequency bands used for both training and testing. The pattern of the matrices is informative regarding whether stimulus information is broadband, narrowband, or if distinct information is contained between the frequency bands. This analysis can be time resolved but is inherently imprecise due to the nature of spectrograms. Therefore, we broadly sampled from representative times [-100ms, 60ms, 100ms, 200ms, 350ms] to capture how frequency information evolves over time.

## Results

### Dissociated information in volume conducted versus locally generated LFP

LFP signals were recorded using linear microelectrode arrays affording laminar localization and alignment (Figure 1C). Next, we recomputed the locally generated LFP – both broadband and in distinct frequency bands – from the volume conducted signal using CSD as an intermediary. This process is detailed for an example session in Figure 3. The volume conduction-containing versus the locally generated, near-field LFP will hereafter be referred to as LFP and nfLFP, respectively. We assured the transformation between LFP and nfLFP was impactful by evaluating the power spectral density (PSD) along the layers of cortex (Figure 3). This analysis demonstrates a difference in spectral power and content between the two LFP signals. However, our main interest was to evaluate the difference in information content between the LFP and nfLFP.

To compare how stimulus responses differed and evolved over time and space for the LFP and nfLFP signals, we extracted information about stimulus features for each of the signals by performing a moving searchlight analysis (Etzel et al., 2013; Tovar et al., 2020). For this analysis, we trained and tested a linear discriminant analysis (LDA) classifier for each session using one electrode contact and its immediate neighboring electrode contacts at each timepoint, iteratively repeating the process until we performed the analysis for the entire laminar array for the interval around stimulus presentation [-100 ms to 400 ms post stimulus onset]. This analysis creates spatiotemporal maps of information flow across the V1 laminar microcircuit for each electrode insertion. To quantify the information that contaminates the LFP signal by volume conduction, we subtracted the information flow maps for nfLFP from those for LFP. For both the LFP and nfLFP information flow maps, we used Wilcoxon signed rank tests to evaluate for statistical significance compared to chance decoding, with FDR correction for multiple comparisons over time and space. Chance decoding is defined as the percentage of correct decoding that is obtained when the classifier guesses randomly. All results were thresholded by statistical significance, following FDR-correction across electrodes and time, at q<0.025. For the LFP and nfLFP differences, significance was evaluated against zero.

For eye-of-origin information (Figure 4A), decoding from LFP and nfLFP differed primarily during the initial transient response. In the LFP, eye-of-origin information first emerged in the granular layer and quickly spread to the supragranular layer, while being greatly diminished in the infragranular layers. Conversely, in the nfLFP signal, information was present in all layers, including the infragranular layers. This result is notable given previous findings of reduced eye-of-origin information in infragranular neural spiking (Dougherty et al., 2019; Hubel and Wiesel, 1972; Tovar et al., 2020). This apparent discrepancy can be understood by nfLFP indexing subthreshold synaptic inputs (Buzsáki et al., 2012; Mitzdorf, 1985) while spiking gives rise to suprathreshold output signals (Leszczyński et al., 2020). Most importantly, there was more decodable information in the nfLFP signal throughout the transient response, as shown by the subtraction of nfLFP from the LFP signal. The most prominent difference occurred in the infragranular layers at the time of the response onset transient and shortly following the transient response. Together, these findings show that the volume conducted signals from surrounding cortical columns and other areas in the brain add noise and thus mask true signals containing eye-of-origin information.

**Figure 4:**
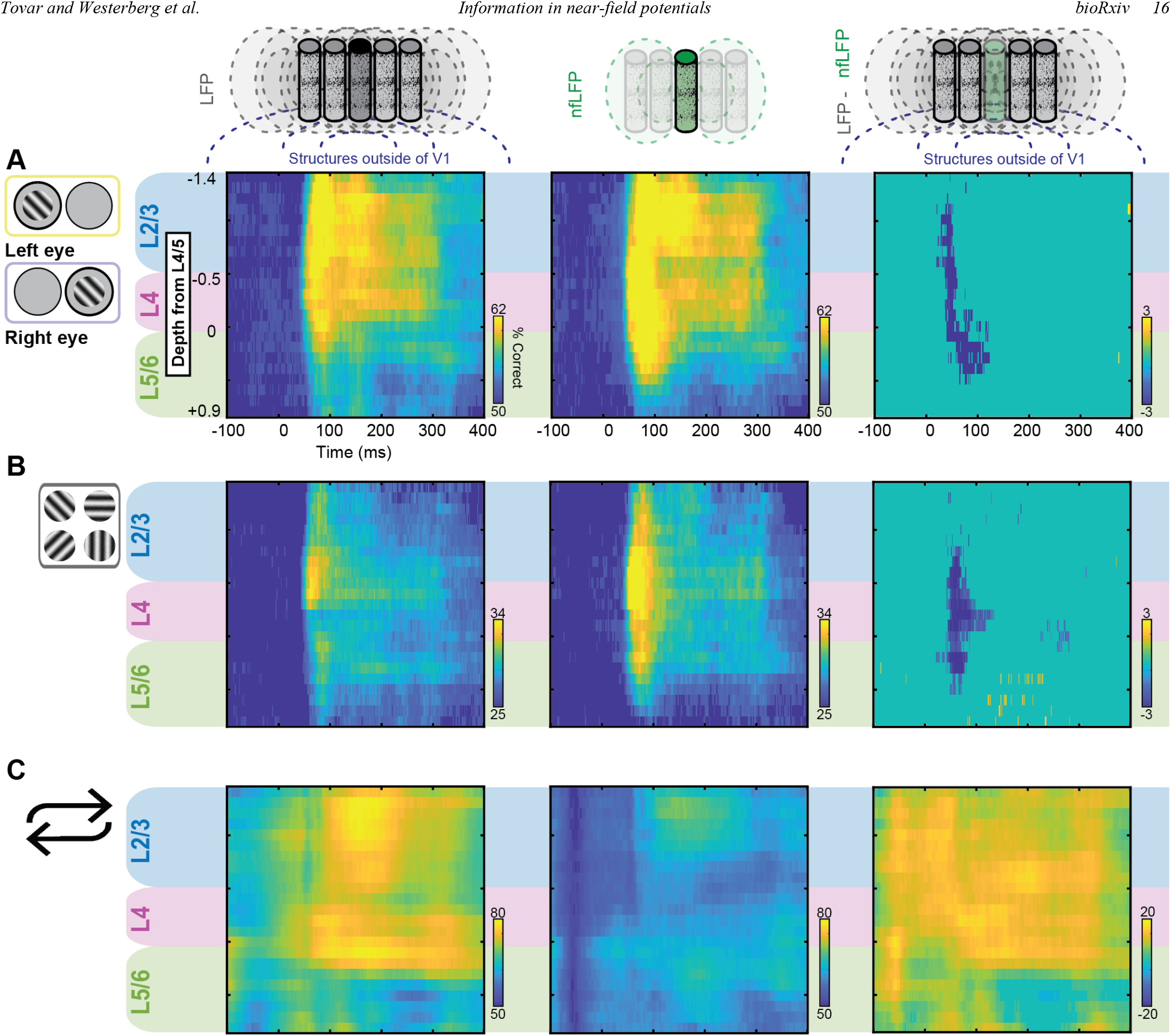
Reducing LFP to its local components (nfLFP) enhances decoding of local processing. A 3-electrode searchlight along the laminar probe was used to decode stimulus (A) eye-of-origin, (B) orientation, and (C) temporal order within repetitions. Left column shows results for the regular LFP signal, middle column shows nfLFP decoding results and right column shows the nfLFP searchlight maps subtracted from the LFP searchlight maps. All results tested for significance against chance decoding: 50% for eye-of-origin and stimulus repetitions, 25% for orientation, and 0% for the searchlight map differences, FDR corrected q=0.05. Ordinates are cortical depth, relative to the L4/5 boundary. Abscissa shows time in milliseconds (0 = stimulus onset). Blue indicates lower decoding performance and yellow higher decoding performance.

The nfLFP signal also contained more orientation information than the LFP signal (Figure 4B), differing most prominently during the transient response. LFP signals carrying information about stimulus orientation spread out unevenly across V1 layers, peaking primarily in the supragranular layers. In stark contrast, the nfLFP transient response carried orientation information evenly throughout the compartments. The orientation information in the nfLFP also appeared to be more prolonged than the LFP. These differences become fully evident when subtracting the two signals, with the most sustained differences located in the granular middle layers. These results show that in addition to eye-of origin, volume conduction obscures information about stimulus orientation in V1 LFP.

However, for stimulation history, this pattern of nfLFP containing more information than LFP was broken (Figure 4C). Using LFP, stimulation history was most reliably decoded from the granular and supragranular layers. However, for the nfLFP signal, information on stimulation history was most prominent in the supragranular layers and infragranular layer. The nfLFP spatiotemporal profile is more consistent with previous reports that relied on spiking responses (Van Kerkoerle et al., 2014; Westerberg et al., 2019). These results show that while stimulus history is represented in V1, this information is found to a greater degree in distant signals that are passively propagated via volume conductions than local LFP. Together, these results show how volume conduction selectively increases or decreases decodable information from cortical LFP.

### Unique frequency patterns convey information regarding stimulus features over time

Next, we investigated how the relative distribution of information regarding stimulus features in different LFP frequency bands is affected by volume conduction. To this end, we devised a novel frequency generalization analysis. We began by constructing a time-frequency spectrogram, collapsed across trials regardless of stimulus features. This time-frequency spectrogram was used to identify key epochs of interest. Specifically, we selected the following epochs (time intervals): (1) baseline prior to stimulus presentation, (2) transient response peak immediately following stimulus presentation, (3) sustained response, (4) stimulus offset, and (5) return to baseline following stimulus presentation. Note that the signal of interest was always centered within each of these windows to compensate for the temporal imprecision that is inherent to spectral decomposition (Figure 5A). In Figure 5B, we show a schematic outlining frequency generalization analysis. Briefly, we trained a classifier at one LFP frequency band and then tested the classifier at all remaining frequency bands to reveal how well different types of stimulus feature information at a particular frequency band generalize across frequency bands. We iteratively repeated this process until we obtained a matrix of all frequency bands used for both training and testing. This analysis was done separately for the selected epochs of interest.

**Figure 5:**
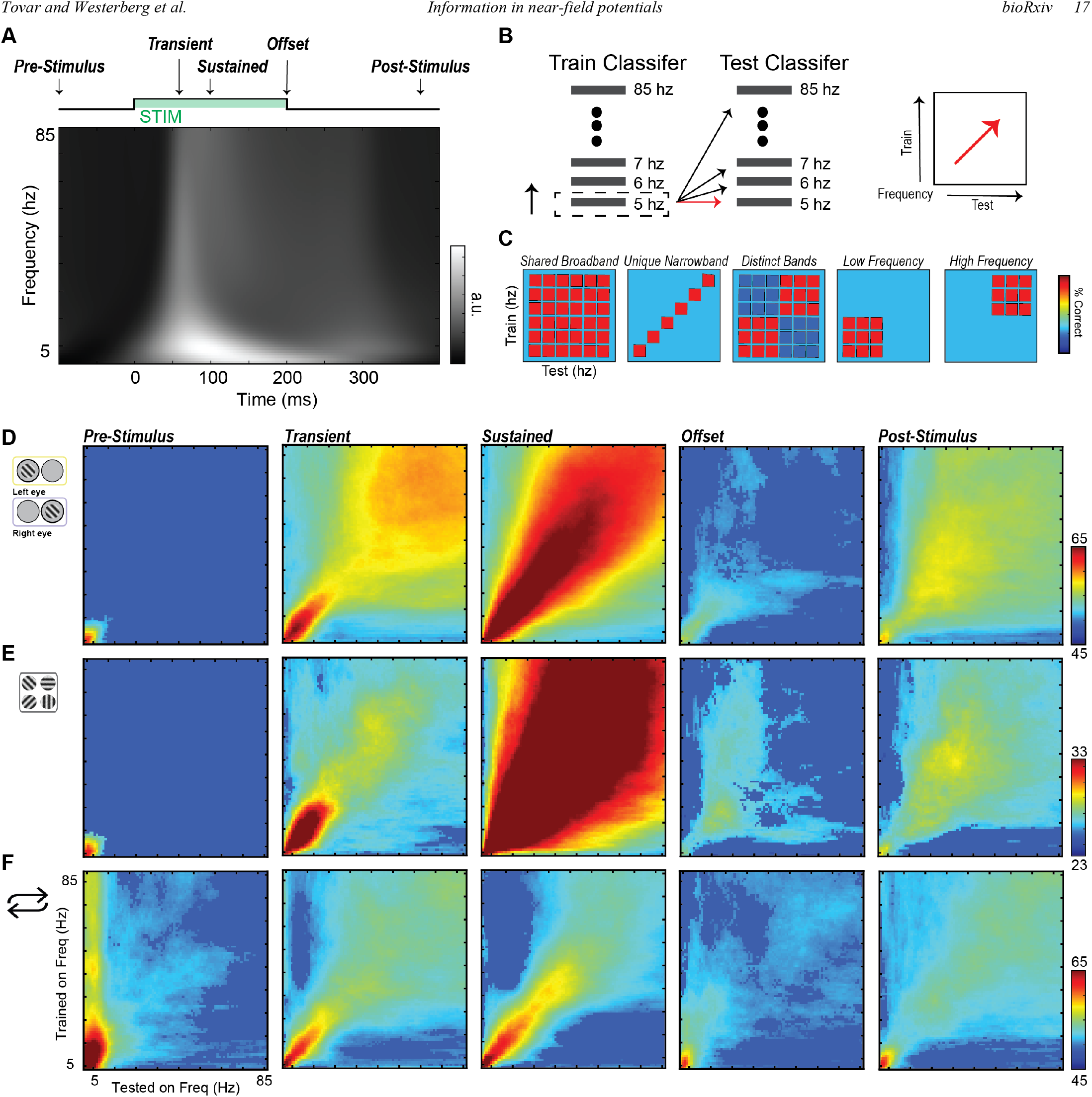
Frequency-dependent generalization of stimulus information evolving over time. (A) Grand average nfLFP full wave rectified time frequency spectrogram of stimulus responses. Stimulation timeline shown above for reference, indicating five key timepoints of interest (B) Schematic of time frequency generalization procedure: An LDA classifier was iteratively trained on a frequency bin and tested on all frequency bins, repeating the process until all frequency bins are used for training and testing to create a frequency generalization matrix. This procedure was done for all five key timepoints of interest. (C) Cartoon models of possible results. (D-F) Frequency generalization matrices FDR corrected for multiple comparisons, q<0.05, for: (D) Eye-of-origin, (E) Orientation, (F) Stimulus repetitions.

In Figure 5C, we show cartoon models for predictions of matrix patterns that could arise from frequency generalization analysis. Each of these patterns has a unique interpretation. For example, if sensory information is broadly shared across LFP frequency bands, we would expect a square like pattern. If, on the other hand, each frequency bands contains unique information, we expect a diagonal pattern as result. Conversely, if information is contained within distinct bands, we would expect two distinct squares to emerge within the generalization matrix. Lastly, we show the expected pattern if information were contained amongst low frequency bands or high frequency bands exclusively. While these are discrete examples, we deemed it most likely that stimulus information is actually represented by complex combinations of these simple patterns.

In Figures 5D-F, we show the frequency generalization matrices at various times for eye-of-origin, orientation, and stimulus history evolving over time. All matrices were thresholded for statistical significance using FDR correction for multiple comparisons at q<0.05. We will break down the differences across our epochs of interest: (1) During the pre-stimulus period, we found some significant decoding for both eye-of-origin and stimulus orientation within the lowest frequency bands. This paradoxical result is most likely due to lack of appropriate temporal resolution. This assumption is supported by the fact that stimulus onset did not alter decoding, indicating “smearing”. For stimulation history, however, we also found significant decoding in high frequency bands before the stimulus was even presented. This result can be explained by changes in baseline activity following stimulus presentation (baseline shifts). In other words, following stimulus offset, the broadband LFP does not seem to fully recover to the initial pre-stimulus baseline, thus allowing for decoding of stimulation history. (2) During the initial, transient response peak, marked differences emerged. Eye-of-origin and orientation both showed a broadband pattern, though orientation information was shared more widely across frequencies. Stimulus history on the other hand, contained distinct low and high frequency bands of information, roughly corresponding to sub-gamma (<30Hz) and high gamma activity. (3) During the sustained response period immediately following the initial response transient, eye-of-origin information could be reliably decoded bispectrally from low and high frequencies, one unique narrowband from 5-25 Hz and a broader band, ranging from 45-85 Hz.m Orientation information on the other hands was largely confined to the lower frequency bands at <25 Hz. The spectral distribution of information about stimulation history did not considerably change from the transient peak. (4) At stimulus offset, stimulus information is reduced across stimulus dimensions with notable reduced performance in the lower higher and lower frequencies for eye of origin information when compared the sustained responses. (5) Following stimulus offset, stimulus feature information dissipates with only some remnants of eye-of-origin and stimulus history information in the low frequency bands (>20 Hz), suggesting an alteration of V1’s state that goes beyond sensory responses in the form of baseline changes. Together, these results demonstrate how feature information contained in different frequency bands vary over the course of a stimulus presentation.

### Information decoded from LFP and nfLFP differs between frequency bands

We next quantified how volume conduction affects different frequency bands of the LFP. Using LFP power and nfLFP power signals (Figure 2C), we again employed a moving searchlight analysis to construct spatiotemporal maps for stimulus decoding. All results were thresholded by statistical significance, following FDR-correction across electrodes and time, at q<0.025. Beginning at theta, a frequency band associated with feedforward activity (Bastos et al., 2015), there was more information in the localized signal in cortical layers associated with the initial feedforward volley (Figure 6A). For eye-of-origin and orientation, information first emerged in the feedforward-associated granular input layers (Casagrande and Boyd, 1996; Hubel and Wiesel, 1972). In contrast, information regarding stimulation history first emerged most prominently in the feedback-associated supragranular layer (Tovar et al., 2020; Westerberg et al., 2019).

**Figure 6:**
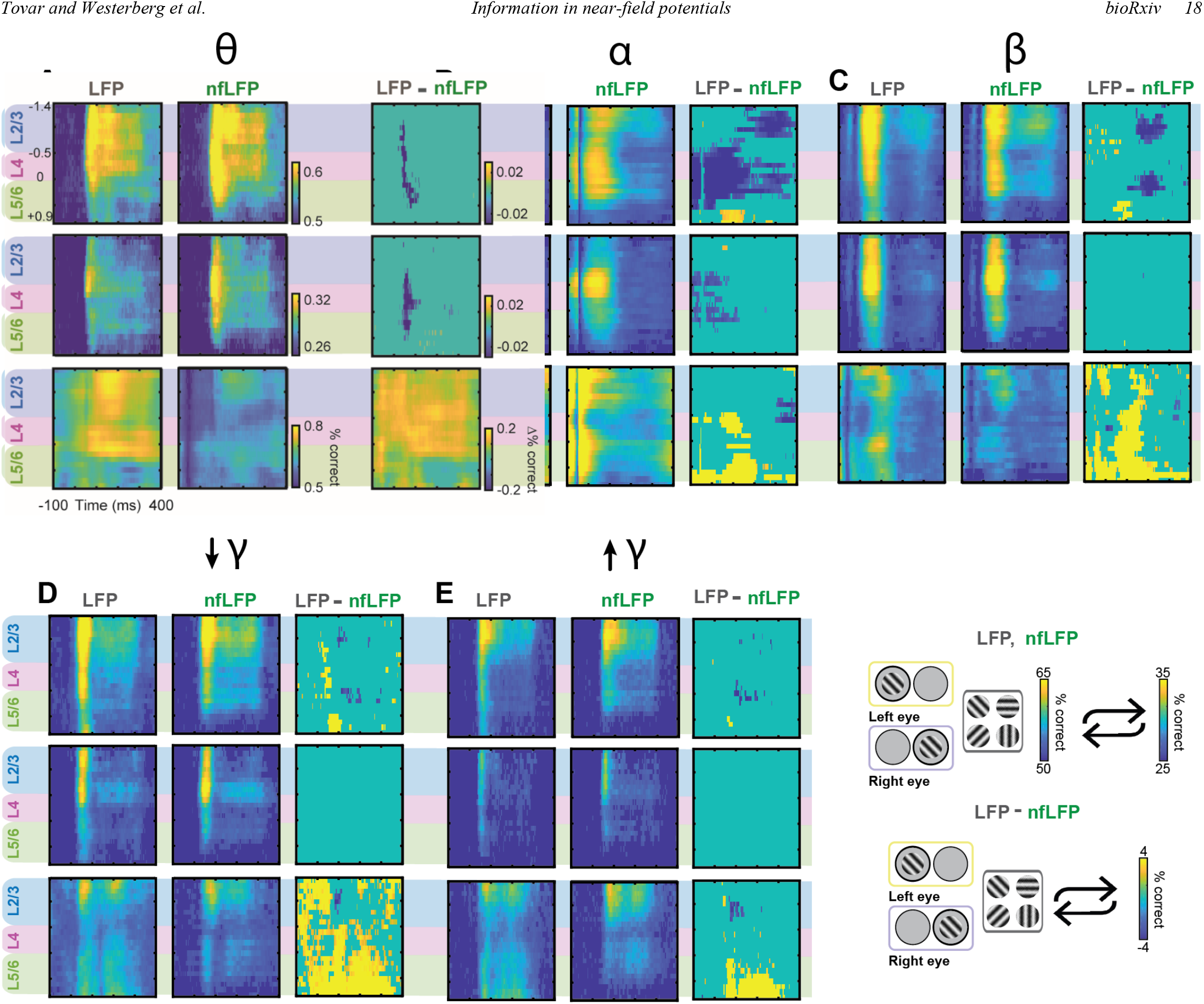
LFP and nfLFP differ across cortical layers and across frequency bands. A 3-electrode searchlight along the laminar probe was used to decode Eye-of-origin, Orientation, and Stimulus Repetitions for (A) theta 4-8 Hz, (B) alpha 8-15 Hz, (C) beta 15-30 Hz, (D) low gamma 30-60 Hz, and (E) high gamma 60-100 Hz. LFP, nfLFP, and difference maps tested for significance from chance decoding, FDR corrected, q=0.05.

During the sustained response, near-field information regarding eye-of-origin and orientation in supragranular and infragranular layers fell off compared to the volume conducted LFP. Meanwhile, there was more information regarding stimulus history decodable from the LFP signal within granular and infragranular layer both before and after stimulus presentation. Given that we found information regarding stimulation history prior to onset of repeated presentations, it is not surprising to find significant decoding at stimulus offset either. In total, we found that in theta, layers that are not associated with the initial volley of sensory information are the layers in which volume conduction effects predominate.

For alpha activity (Figure 6B), there was more eye-of-origin information in the localized nfLFP signal both for the initial feedforward sweep in the granular layers as well as the supragranular layers for the sustained response. These results are consistent with recent work (Gieselmann and Thiele, 2020) demonstrating that alpha is associated with feedforward inputs in addition to the more commonly associated feedback processing (Buffalo et al., 2011; Van Kerkoerle et al., 2014). By removing volume conduction, eye-of-origin information in the feedforward sweep becomes more apparent. For stimulus orientation, there was more localized information along the transient, spread out equally across layers during the initial feedforward sweep. These results suggest that the volume conducted signal adds noise to stimulus orientation information, as decoding improves for the reduced nfLFP signal. For stimulation history, the LFP signal predominated throughout the spatiotemporal map, suggesting that information regarding stimulus repetition primarily originates outside V1. Overall, these results demonstrate the utility of the nfLFP in highlighting the role alpha may have in feedforward processing for select stimuli features.

In the beta range (Figure 6C), the nfLFP signal from the supragranular and infragranular layers near the 4/5 border contained more eye-of-origin information during the sustained response, mirroring alpha activity. Similarly, but to a much lesser extent, low gamma and high gamma eye-of-origin information showed differences during the sustained response (Figure 6D-E). Low gamma, but not high gamma activity showed more information in the LFP signal than the nfLFP signal, suggesting that high gamma power is more reflective of local processing. Meanwhile, differences in orientation information between the LFP and nfLFP were minimal, ranging from beta to high gamma (Figure 6C-F). For stimulation history, there was more information in the LFP signal than nfLFP signal throughout beta to high gamma frequencies (Figure 6D-E). Interestingly, this information became increasingly localized to the infragranular layer at higher frequencies. These results are consistent with previous studies that found that volume conduction effects are most prominent in the infragranular layers (Kajikawa and Schroeder, 2015). Overall, across frequencies, different spatiotemporal profiles emerged for the LFP signal and CSD-derived nfLFP signal, depending on the stimulus property, suggesting different degrees of volume conduction.

### nfLFP is more informative than spiking regarding local cortical stimulus processing

Lastly, we compared the amount of information content and its spectral and laminar distribution between nfLFP and population spiking (multiunit activity, MUA). This is both an interesting question from a theoretical point of view as well as an important question from a practical perspective.

To address this question, we contrasted the decoding accuracy across space in time for each stimulus feature between nfLFP and population spiking (MUA) (Figure 7). To our surprise we found that nfLFP is significantly more informative with respect to local computations, as it yields greater decoding accuracy across space and time than population spiking. Remarkably, this was even true, albeit to a much lesser degree, for stimulus repetition, which likely has a non-local origin. Thus, LFP, once it has been computationally de-artifacted from volume conduction, has more decoding power than columnar spiking activity. This superiority encompasses both local computations as well as (likely) feedback-associated processes, perhaps due to the fact that LFP is more sensitive to (subthreshold) synaptic inputs and processing than (suprathreshold) spiking outputs. When it comes to maximizing the read-out of informative brain activity, nfLFP therefore seem to be the signal of choice, at least when columnar cortical processing is concerned.

**Figure 7:**
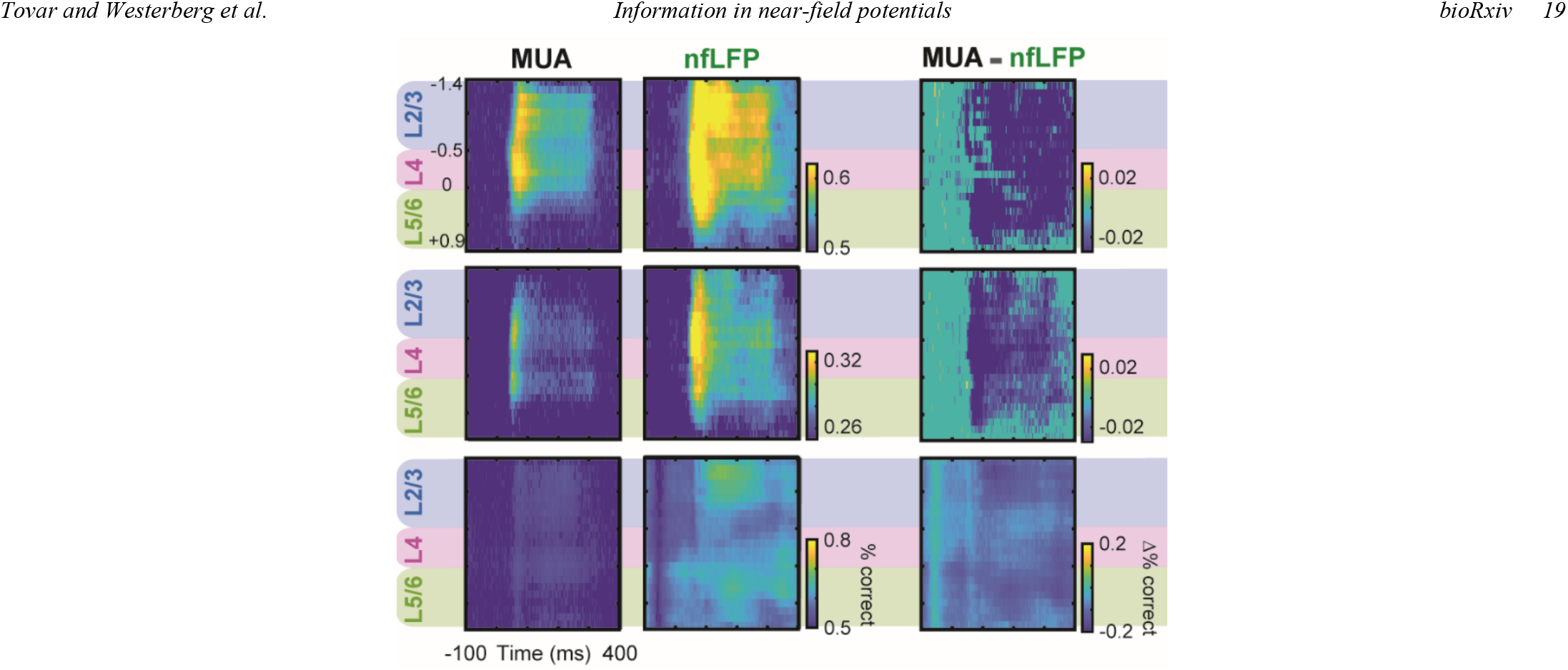
Relative information in nfLFP and population spiking (MUA) across cortical layers. Left column: MUA. Middle column: nfLFP. Right column: Subtraction (shades of blue indicate that nfLFP was more informative than MUA). Top row: Stimulus eye-of-origin information. Middle row: Stimulus orientation information. Bottom row: Stimulus history (repetition) information. Note that for all three types of stimulus information, nfLFP decoding yielded better classifier performance, suggesting that the information content in nfLFP is higher than that contained in population spiking.

## Discussion

We used MVPA to extract information regarding various stimulus properties from regular, volume conduction (far field)-contaminated LFP and the localized, near-field nfLFP signal. Information was defined - and quantified - as the ability of a classifier to extract meaningful information regarding both concrete (e.g., physical makeup) and abstract (e.g., historical context) properties of sensory stimuli. By analyzing more than just response magnitude of these respective signals, we were able to show that LFP components that arise by volume conduction reduce information about local neural processes. These far-field contributions also enhance information about (presumably) non-local neural processes, which could cause these responses to be mistaken as of local origin. This finding is remarkable in its counter-intuitive nature: The process of stripping LFP of volume-conducted components (i.e., double differentiation) increases random noise fluctuations that are inherent in the data, and thus reduces the overall signal-to-noise ratio (SNR). Yet, despite this reduction in overall SNR, decodable information about locally performed computations *increased*. We further explored how sensory information varies as a function of LFP frequency. This analysis, too, showed that volume conduction can lead to erroneous attribution of functional roles for certain frequency bands, since the nfLFP contains information in different frequency bands than regular LFP. Together, these results provide an analytical framework that can be applied to any brain area to decipher whether feature information is contained to a local circuit or relayed from remote sources.

It is important to note that our findings have much more wide reaching implications than studying different neural signals and the capability of ML-based decoding. Intracranial implants are increasingly used to both read and write brain activity in human patients. The promise of brain-machine interfacing (BMI) to e.g., control robotic appliances by measuring and decoding intracranial cortical activity is particularly exciting. Our work suggests that these techniques would benefit greatly from not only decoding regular, far-field contaminated LFP and spiking, but also consider near-field LFP or CSD. The obvious challenge that comes with this approach is that it requires high electrode density within a limited volume. However, the ongoing trend of using arrays for BMI already affords this possibility. All that would then be needed is an added analytical or data processing step involving differentiation between neighboring electrodes to eliminate shared (likely volume-conducted) signals.

There is longstanding debate about the spatial extent of the local field potential (Kajikawa and Schroeder, 2015, 2011; Katzner et al., 2009; Mineault et al., 2013; Xing et al., 2009). Our results suggest that the answer to this question might be best answered by considering information content. Both experimental and modeling studies have shown that specific properties of the LFP signal, primarily correlated synaptic inputs, can differ by an order of magnitude in volume conduction (Buffalo et al., 2011; Leski et al., 2013; Lindén et al., 2011; Rosenbaum et al., 2017). Not surprisingly, one of the biggest factors that influences the correlation of synaptic inputs is the information content of the stimulus being processed (Peter et al., 2019).

The debate regarding the extent of volume conduction also extends into frequency space. Here, the question is whether volume conduction effects occurs evenly across frequencies (Kajikawa and Schroeder, 2015, 2011), or whether volume conduction is more prominent for some frequency bands than others (Leski et al., 2013). Much like the broader question regarding broadband LFP spread, apparent discrepancies between studies might be explained contextually by the specific information contained within different frequency bands. Differences between frequency bands have been highlighted by a number of studies investigating feedforward and feedback activity (Bastos et al., 2015; Van Kerkoerle et al., 2014), signal synchrony along frequencies (Buffalo et al., 2011), or shared mutual information using information theoretic approaches (Belitski et al., 2008; Kayser et al., 2009). However, contextual parameters such as stimulus size or attentional selection can influence synchronization within frequency bands (Buffalo et al., 2011; Ferro et al., 2021; Gieselmann and Thiele, 2020). In the current study, we show that within stimulus presentations, shared information between frequency bands changes radically depending on both the stimulus feature and the particular time epoch of the neural response.

A valid question is why we quantified information content in LFP at all. Since volume conduction contaminates information, why did we not stick to CSD or nfLFP? Our study shows that there can be significant added value in determining the information content of volume conducted signals. For example, we found that stimulus history information was more prominently found outside than within V1 microcircuits. This finding is consistent with what is known about the role of many brain areas including the visual pulvinar, middle and inferior temporal gyri, and frontal gyri in repetition suppression (Kaas and Lyon, 2007; Wig et al., 2009). However, it is also possible that this contrast can lead to new discoveries. Recently, convolutional neural networks (CNNs) have been introduced to model ventral visual stream with remarkable accuracy (Kar et al., 2019; Kar and DiCarlo, 2020; Schrimpf et al., 2018; Yamins and DiCarlo, 2016). However, while we can visualize features captured by CNN layers (Bashivan et al., 2019), these features are not anchored in cognitive theories or experiments specifically targeting those features. In this circumstance, contrasting local signals to volume conducted signals could help decide whether a particular CNN layer matches local computations, or are likely to be found outside of the recording site.

There are also at least two important theoretical consideration surrounds a comparison of decoding spikes versus nfLFP. First, from a theoretical point of view, there has been the long-standing observation that LFP are fundamentally different from local spiking (Logothetis et al., 2001). This difference could be explained by LFP representing synaptic “inputs” that contrast with spiking “outputs” in that the difference between them constitutes local computations. However, due to the fact that LFP is generally contaminated with non-local, volume conducted signals, computing this contrast has generally proved difficult. By contrasting nfLFP with spiking, we can get closer to this idealized experiment.

From a practical point of view, there is a perhaps more pressing question as to which neural signal is best selected to “read out” brain activity in the context of a brain machine interface (Andersen et al., 2004). This is a multifaceted question since there are various pros and cons for various signals that are independent of maximized information content (such as required invasiveness or temporal stability). However, at least to our knowledge, it is largely unclear which neural population signal maximizes decoding from a limited amount of brain tissue. In particular, it has long been unclear whether LFP is more or less informative than neuronal spiking once volume conduction has been taken into account. Our results provide a clear - and highly surprising - answer to this conundrum in that spiking activity alone may not be the most desirable measure for brain-machine interfaces.

To sum, we have demonstrated utility of using MVPA to extract feature specific information from local and distant signals in the V1 microcircuit. The different spatiotemporal profiles between LFP and nfLFP for eye-of-origin, orientation, and stimulus history highlights the importance of accounting for possible contamination from distant signals. By focusing on stimulus features, rather than activation, our results also help reconcile conflicting findings from previous studies quantifying volume conduction in the LFP signal as a whole, as well as how volume conduction differs by frequency. Lastly, our study provides a method with potential practical applications. For example, if it were only possible to record from an early cortical area and late cortical area, but not an intermediate area, comparing between LFP and nfLFP could provide clues on how information transforms between these two areas. This added flexibility can be invaluable in understanding cognitive processes when the number of recording sites is limited by practical or theoretical constraints.

## Acknowledgements

This work was supported by NEI (R01EY027402 and P30EY008126) and the NIH Office of the Director (S10OD021771). D.A.T. was supported by a training grant from NIGMS (T32GM007347). J.A.W. was supported by fellowships from NEI (F31EY031293 and T32EY007135). A.M.B. was supported by NIMH grant 4R00MH116100-03. The authors would like to thank M. Feurtado, I. Haniff, M. Maddox, S. Motorny, D. Richardson, M. Schall, L. Toy, B. Williams, and R. Williams for technical support.

